# Comparison of two commercial ELISA kits for detection of rubella specific IgM and IgG antibodies

**DOI:** 10.1101/305813

**Authors:** Rajlakshmi Viswanathan, Suji George, Manoj V Murhekar, Asha Mary Abraham, Mini P Singh, Santoshkumar M Jadhav, Vijayalakshmi Nag, Sadanand Naik, Chandrashekhar Raut, M Ashok, Minakshi Gupta, Vishal Jagtap, Ojas Kaduskar, Nivedita Gupta, Gajanan N Sapkal

**Affiliations:** Diagnostic Virology Group, Indian Council of Medical Research-National Institute of Virology, Pune, India; Indian Council of Medical Research-National Institute of Epidemiology, Chennai, India; Department of Clinical Virology, Christian Medical College, Vellore, India; Department of Virology, Post Graduate Institute of Medical Education & Research, Chandigarh, India; Bioinformatics and Data Management Group, Indian Council of Medical Research-National Institute of Virology, Pune, India; Department of Microbiology, All India Institute of Medical Sciences, Jodhpur, India; Department of Biochemistry, King Edward Memorial Hospital, Pune, India; Indian Council of Medical Research-National Institute of Virology, Bengaluru Unit, India; Department of Microbiology, Tata Main Hospital, Jamshedpur, India; Division of Epidemiology and Communicable Disease, Indian Council of Medical Research, New Delhi

## Abstract

Enzyme linked immunosorbent assay (ELISA) plays an important role in laboratory confirmation of congenital rubella syndrome (CRS), postnatal rubella and seroprevalence studies in different populations. Variation of results are documented for samples tested by different commercial kits. The Enzygnost rubella ELISA, widely used in the WHO network, is expensive and not readily available. In the present study, performance of the Euroimmun ELISA was compared to the Enzygnost ELISA for detection of rubella specific IgM and IgG antibodies.

Two hundred and eighty five serum samples collected from suspected CRS patients identified through a recently initiated surveillance for CRS at six sentinel hospitals and 435 serum samples from a serosurvey of pregnant women from these sites, were available for testing of rubella specific IgM and IgG antibodies respectively. Qualitative agreement (concordance percentage and Cohen’s Kappa coefficient -κ) was evaluated for both IgM and IgG assays. Bland – Altman plots were used to assess the difference in quantitative agreement for IgG titers.

Good qualitative agreement between the two ELISA kits was observed for detection of both anti rubella IgM (94.7% agreement and κ of 0.86) and IgG (96.3% agreement and κ of 0.84). Sensitivity and specificity of Euroimmun assays compared to Enzygnost was 100% and 93.1% for IgM and 95.9% and 100% for IgG respectively. Bland – Altman analysis for paired quantitative results of rubella specific IgG yielded a mean difference of 0.781 IU/ml with majority of values (97.1%) within ± 2 SD of the mean difference. Euroimmun ELISA provided on an average, higher titers as compared to Enzygnost.

Our study findings suggest that Euroimmun ELISA may be considered for detection of rubella specific IgM in suspected CRS cases and rubella specific IgG in surveillance studies.

## Introduction

Rubella is an acute, contagious vaccine preventable viral infection, which usually causes a mild exanthematous febrile illness in children or young adults. Infection with rubella virus during pregnancy, particularly in the first trimester, can cause congenital rubella syndrome (CRS), which may result in malformations, disability and even death of the fetus. In September 2013, the WHO Regional Committee for South-East Asia decided to adopt the goal of measles elimination and rubella/CRS control in the South-East Asia Region by 2020 (1). The Government of India has made a commitment towards measles elimination and rubella and CRS control by 2020. Laboratory based surveillance for measles-and rubella is well established in India. Surveillance for CRS has been recently been initiated in six sentinel sites in 2016 (Murhekar M et al. Sentinel surveillance for congenital rubella syndrome — India, 2016-2017. Submitted for publication). In order to estimate rubella sero-prevalence among pregnant women, the first serosurvey was conducted between March/April – May/June 2017, at these sentinel sites (Muliyil D et al. Sero-prevalence of rubella among pregnant women in India, 2017. Submitted for publication).

Enzyme linked immunosorbent assay (ELISA) plays an important role in laboratory confirmation of congenital rubella syndrome and post natal rubella (2). Laboratory tests recommended for confirmation of CRS include detection of rubella specific IgM antibodies in younger infants (<6 months of age) and sustained rise of rubella specific IgG antibodies in older infants (≥ 6 months) (2). In addition, detection of rubella specific IgG antibodies is important to estimate the seroprevalence in different populations, including pregnant women (3,4).

In the facility based surveillance for CRS, a commercial ELISA kit (Euroimmun, Luebeck, Germany) is used for laboratory diagnosis, after considering factors like ease of procurement, timeliness of supply, experience of laboratory experts and cost. The performance of Euroimmun ELISA was evaluated against another reference ELISA kit- the Enzygnost Rubella ELISA kit (Siemens Healthcare GmbH, Henkestr. Erlangen, Germany). This kit is widely used across the WHO measles-rubella network, including in India. Comparability of results was studied, for detection of rubella specific IgM and IgG antibodies, among suspected cases of CRS and pregnant women respectively.

## Materials and Methods

### Ethics statement

The study was approved by the Institutional Ethics committees of all sentinel sites. Participants were enrolled in the study after obtaining written informed consent from pregnant women and parents / guardians of infants.

### Study population and sample selection

Suspected CRS cases were identified among infants (0-11 months) attending pediatrics, ear nose and throat (ENT), ophthalmology, and cardiology Outpatient Departments (OPDs) of six sentinel hospitals. These are tertiary care centers providing medical care to local children as well as those from surrounding states. Pregnant women attending antenatal clinics at these hospitals during March/April – May/June 2017 were eligible to participate in the serosurvey.

### Study samples and case definition

Suspected cases were recruited as per WHO-recommended standards for CRS surveillance (5). Serum samples (n=291) were collected, aliquoted and stored (testing aliquot at 2-8 degree C and remaining aliquots at −20°C) until transported in cold chain to the apex laboratory (ICMR-National Institute of Virology, Pune). Procurement of kits was centralized and all the sentinel sites were provided kits transported in cold chain.

### Testing of samples

Testing was performed using commercial ELISA kit for detection of anti rubella IgM and IgG antibodies (Euroimmun, Luebeck, Germany). All the samples from suspected CRS cases were tested for rubella specific IgM antibodies at the sentinel sites. Serum samples collected from 1800 pregnant women in a serosurvey between March/April-May/June 2017, were tested for rubella specific IgG antibodies at the sentinel sites.

At ICMR-NIV, Pune, 285 of 291 serum samples collected from suspected CRS patients and 435 of 1800 samples from pregnant women were available for testing. These samples were tested by Euroimmun ELISA for the purpose of quality control. To determine agreement between the two kits for detection of anti rubella IgM and IgG antibodies, these samples were retested by Enzygnost Rubella ELISA. Anti rubella IgG titres were extrapolated from a four-point standard curve comprising of calibrators with different concentrations of antibodies, as per manufacturer’s instructions. Titers above the upper limit of detection of 200 IU/ml were denoted as 201 IU/ml and values below lower limit of detection(1IU/ml), were taken as 0.5 IU/ml (Euroimmun). Titres for IgG obtained with the Enzygnost kit, were calculated using the α and β constants whose values are provided with the kit.

Samples, whose results were discordant between the two kits, were retested in duplicate and 2 of 3 results were considered final. All equivocal samples were also re tested in duplicate and any samples which still remained equivocal were excluded from analysis. Chemiluminiscence assay (CLIA) was performed at two of the network laboratories for samples which remained discordant after testing in triplicate. The VITROS 3600 Immunodiagnostic system (Orthoclinical Diagnostics, Raritan, NJ) and the Liaison (DiaSorin, Saluggia, Italy) systems were used for testing IgM and IgG respectively. A sample was considered a true positive or a true negative if concordance of at least two of the three methods was obtained.

### Measurement of intra assay and inter assay precision

The intra assay precision was calculated by testing duplicates of samples within a single run, by the two ELISA systems for both rubella specific IgM and IgG. For determination of inter assay precision, positive controls of same lot provided by the manufacturer, were run repeatedly on different days. The coefficient of variation (CV) was calculated.

The mean absorbance of positive control wells was compared between the two kits for rubella specific IgM and IgG. The characteristics of the kits used in the study are provided (**Supplementary Table1**).

### Statistical Analysis

Qualitative agreement between the two kits, was assessed for both rubella specific IgM and IgG by calculating percent agreement and Cohen’s kappa coefficient (k) statistic to assess the degree of (inter-rater) agreement for serum antibody status. Standard formulae were used to calculate percent agreement, sensitivity and specificity of results by Euroimmun kit in comparison to Enzygnost. As titers estimated by both IgG kits were not normally distributed (Kolmogorov-Smirnov Z=3.15 for Enzygnost and 4.81 for Euroimmun kit), a logarithmic transformation of original data was performed. Paired quantitative measurements of rubella specific IgG (n=422) were then compared by Bland – Altman analysis and plots (6J. Statistical analysis was performed using Open Epi version 3 software and Microsoft Excel.

## Results

### Comparative performance of Euroimmun and Enzygnost kits for detection of Rubella specific IgM

Around 28% (80/283) of the sera from suspected CRS patients were positive by Euroimmun ELISA and 22.9% (65/283) by Enzygnost (*2 samples which gave equivocal results in one or both tests excluded*). A concordance of 94.7% (268/283) was observed for rubella IgM results tested by Euroimmun and Enzygnost assays. Cohen’s kappa (κ) was 0.86 (95% CI: 0.75-0.98), with a sensitivity of 100% (95% CI: 94.4 – 100) and specificity of 93.1.% (95% CI: 88.96 – 95.79) on comparing Euroimmun with Enzygnost results (Table1). Of fifteen discordant samples, nine were of sufficient quantity to be tested by CLIA for anti rubella IgM and results are provided (Table 2).

**Table1:** Qualitative comparison of Euroimmun vs Enzygnost IgM and IgG.

|  |  | Enzygnost IgM ELISA |  |  |
| --- | --- | --- | --- | --- |
|  |  | Positive | Negative | Total |
| Euroimmun | Positive | 65 | 15 | 80 |
| IgM ELISA | Negative | 0 | 203 | 203 |
| Total |  | 65 | 218 | 283 |
| Percent agreement: 94.7%, Kappa: 0.86, Sensitivity: 100%, Specificity: 93.1% |  |  |  |  |
|  |  | Enzygnost IgG ELISA |  |  |
|  |  | Positive | Negative | Total |
| Euroimmun | Positive | 371 | 0 | 371 |
| IgG ELISA | Negative | 16 | 48 | 64 |
| Total |  | 387 | 48 | 435 |
| Percent agreement: 96.3%, Kappa:0.84,Sensitivity:95.9%,Specificity:100% |  |  |  |  |

**Table2:** Comparative analysis of discordant results of rubella specific IgM.

| ID Number | Euroimmun | Enzygnost | CLIA | Final Result |
| --- | --- | --- | --- | --- |
| 1. | Positive | Negative | Non reactive | Negative |
| 2. | Positive | Negative | Non reactive | Negative |
| 3. | Positive | Negative | Non reactive | Negative |
| 4. | Positive | Negative | Non reactive | Negative |
| 5. | Positive | Negative | Positive | Positive |
| 6. | Positive | Negative | Positive | Positive |
| 7. | Positive | Negative | Borderline(1.04)* | Indeterminate |
| 8. | Positive | Negative | Non reactive | Negative |
| 9. | Positive | Negative | Reactive | Positive |

*\* If borderline result is considered positive as the value is close to upper cut off of 1.1,final result may be reported as positive*

The mean absorbance of positive control (PC) wells for 14 assays, by Euroimmun (1.38) was significantly higher (p=0.0009) than Enzygnost PC, for 16 assays (1.02). Good reproducibility was observed for both ELISA systems within the same assay with an average CV of 5.27 for Euroimmun and 6.50 for Enzygnost assays (Table 3).

**Table 3:** Intra assay Precision

| Test | ELISA Kit | Average CV(%) |
| --- | --- | --- |
| Rubella Specific IgM | Euroimmun (n=22) | 5.27 |
|  | Enzygnost (n=18) | 6.50 |
| Rubella Specific IgG | Euroimmun (n=32) | 5.76 |
|  | Enzygnost (n=23) | 6.99 |

**Table 4:** Inter Assay Precision.

| Test | ELISA Kit | Mean(±SD) of Positive control | CV(%) |
| --- | --- | --- | --- |
| Rubella Specific IgM | Euroimmun (n=6) | 1.53± 0.12 | 8.46 |
| | Enzygnost (n=6) | 0.49 ± 0.08 | 16.23 |
| Rubella Specific IgG | Euroimmun (n=4) | 1.44± 0.18 | 12.45 |
| | Enzygnost (n=4) | 1.35± 0.24 | 17.5 |

### Evaluation of the performance of Euroimmun and Enzygnost kits for detection of Rubella specific IgG

Test results of 435 samples were considered for estimation of percent agreement of rubella specific IgG by Euroimmun and Enzygnost kits. Three hundred and seventy one (85.3%) samples were positive by Euroimmun and 387 (89%) by Enzygnost assays respectively. Concordance of 96.3% (419/435) for Euroimmun vs. Enzygnost results was noted. Cohen’s kappa of 0.84 (95% CI: 0.75 – 0.93) was observed, with sensitivity of 95.9% (95% CI: 93.4-97.4) and specificity of 100% (95% CI: 92.6 – 100) on comparing Euroimmun with Enzygnost results (Table1). Of 16 discordant samples, five sera were available in sufficient quantity and tested by CLIA for anti rubella IgG. Results are provided (Table 5).

**Table 5:**
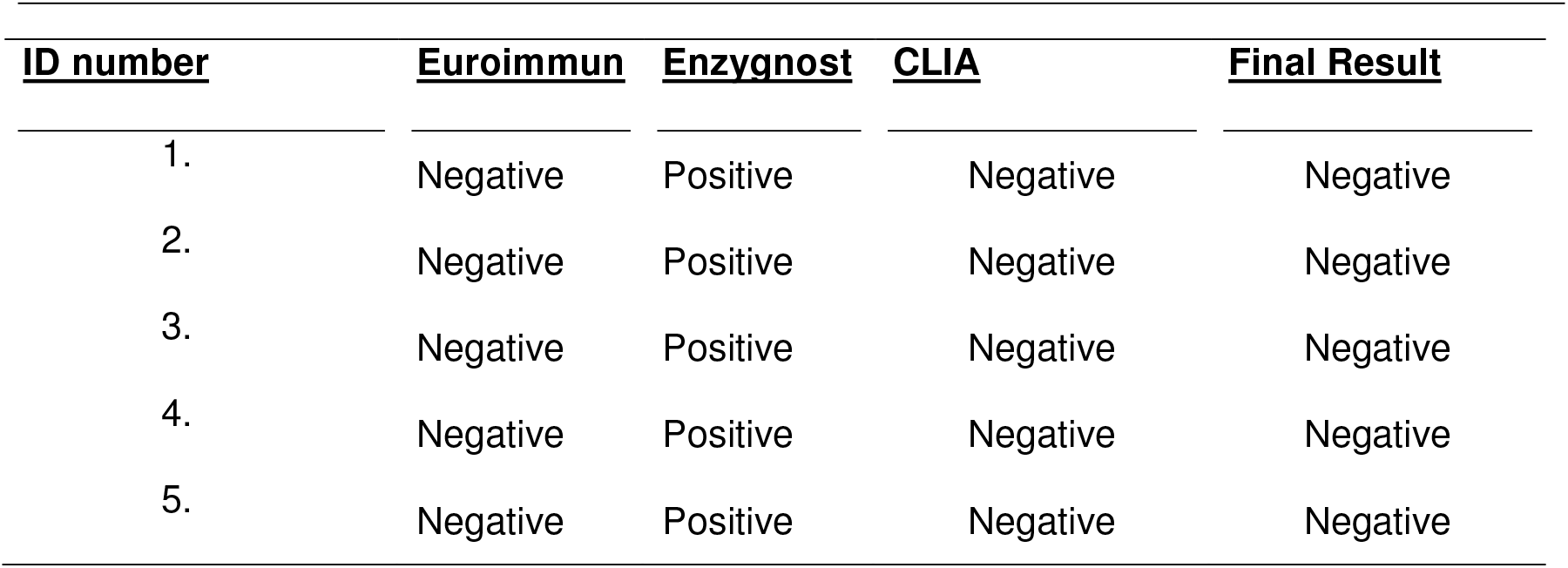
Comparison of anti-Rubella IgG discordance in three systems.

| <u>ID number</u> | <u>Euroimmun</u> | <u>Enzygnost</u> | <u>CLIA</u> | <u>Final Result</u> |
| --- | --- | --- | --- | --- |
| 1. | Negative | Positive | Negative | Negative |
| 2. | Negative | Positive | Negative | Negative |
| 3. | Negative | Positive | Negative | Negative |
| 4. | Negative | Positive | Negative | Negative |
| 5. | Negative | Positive | Negative | Negative |

The mean absorbance of positive control (1.36) of 14 ELISA assays by Euroimmun, was significantly higher (p=0.0025) than Enzygnost positive control absorbance over 16 assays (1.045). Good reproducibility was observed for both ELISA systems within same assay with average CV of 5.76 for Euroimmun and 6.99 for Enzygnost assays (Table 3).

Bland – Altman analysis (Fig1) for paired quantitative results of rubella specific IgG yielded a mean difference of 0.781 IU/ml (95% limits of agreement, 0.725 and 0.840 IU/ml). The majority of values (97.1%) were within ± 2 SD of the mean difference. Euroimmun ELISA provided on an average, higher titres as compared to Enzygnost (Fig 2).

**Figure1:**
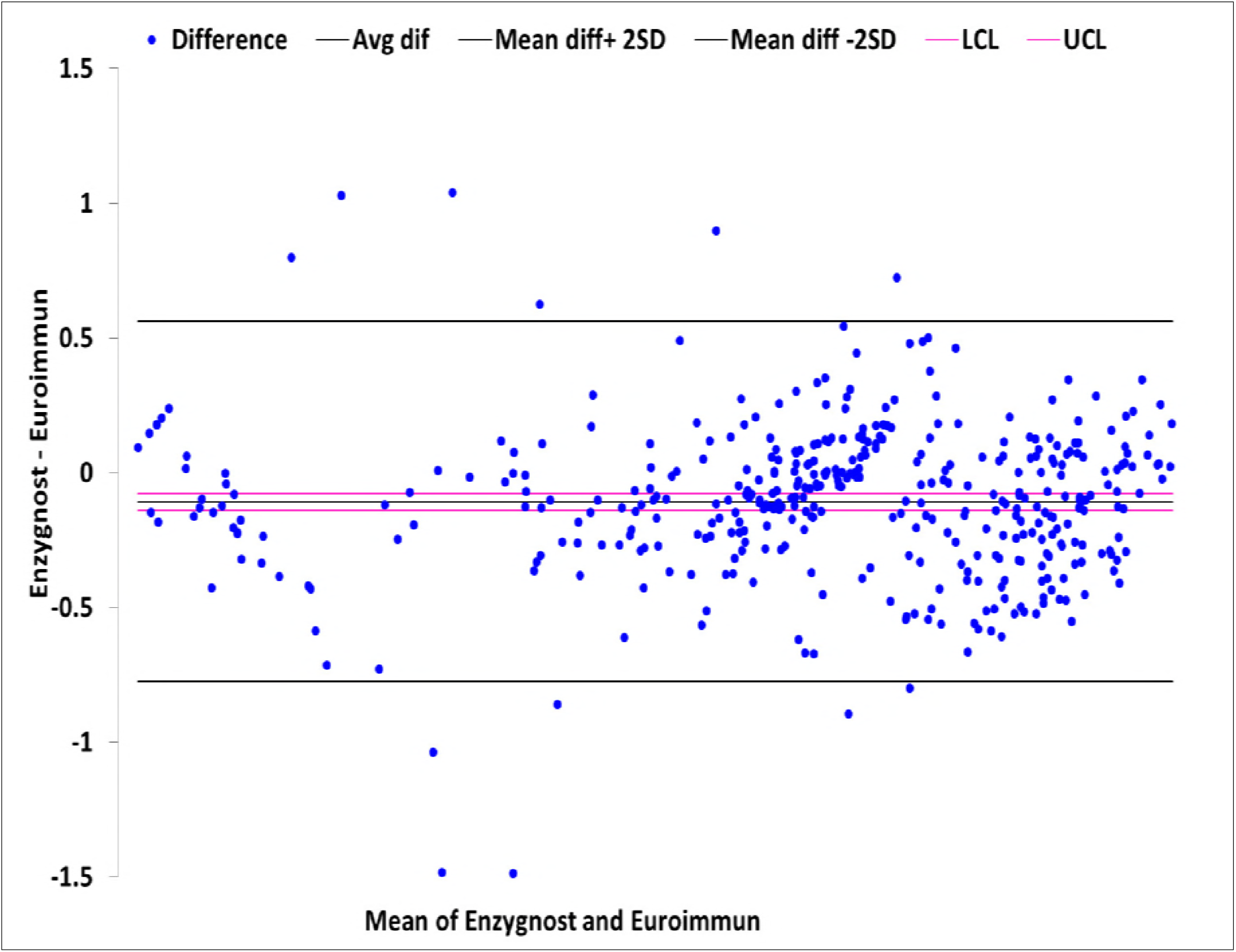
Bland Altman plot showing quantification of agreement between two Methods, IgG titres transformed to logarithm scale.

**Figure2:**
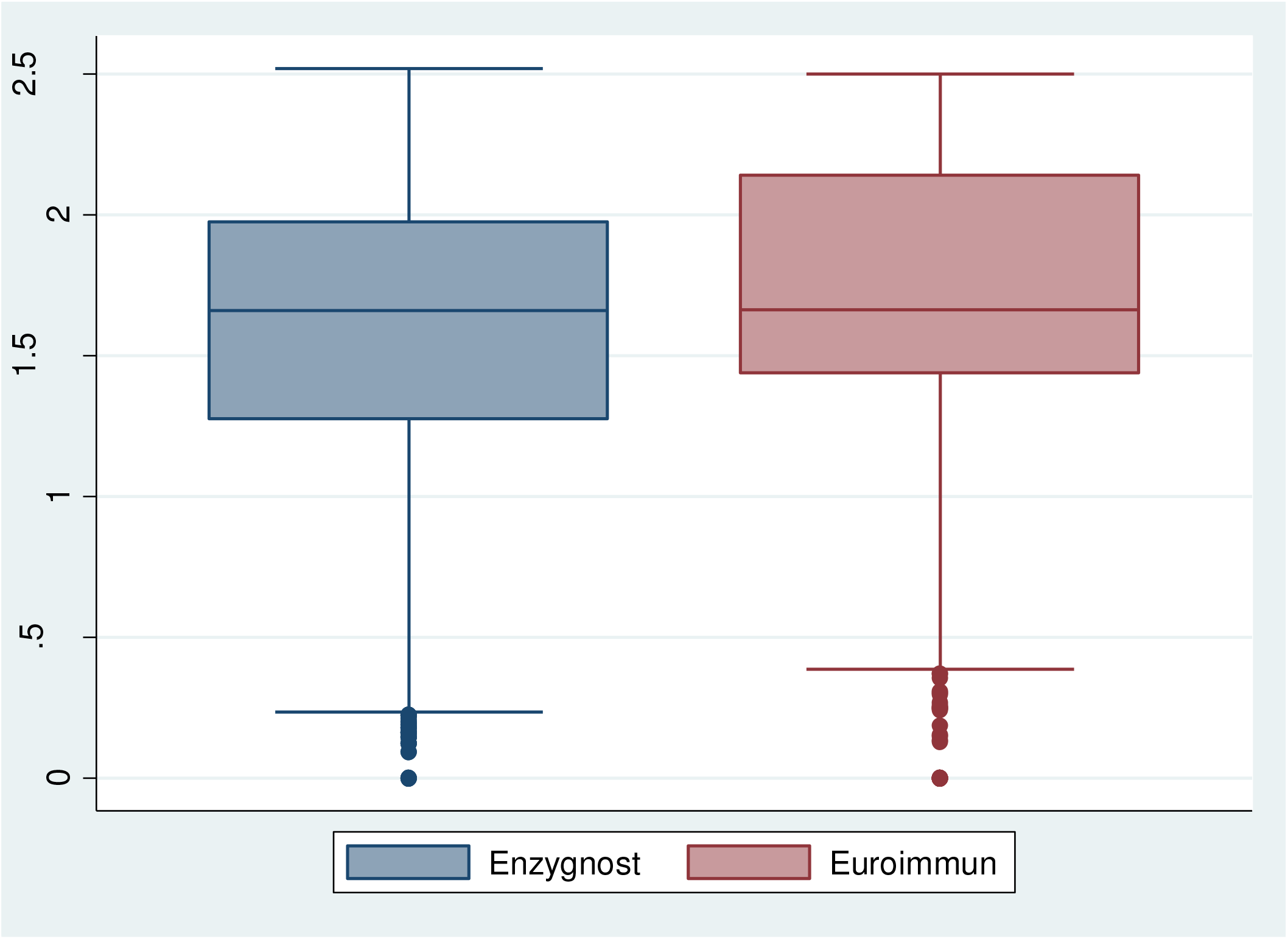
Box plot distribution of IgG titers (Log values-y axis)) from two methods(x axis)

## Discussion

Serological studies of rubella are gaining importance in the light of the ongoing control program for CRS. Detection of rubella specific IgM in suspected CRS cases remains the frontline diagnostic tool for laboratory confirmation and ELISA is one of the most common methods in use. Detection of rubella specific IgG by ELISA, is of value in population based studies, particularly among pregnant women, to understand their susceptibility to infection. The Enzygnost ELISA kit, which is widely used in the WHO network, is also the kit of choice for case based surveillance of measles-rubella in India. Although a well established and validated kit, it is expensive (**Supplementary Table1**) and often unavailable in many laboratories in India, which are not directly linked to the network or which are working on population based studies. This therefore requires the use of other commercial kits, causing variations in reporting of results.

Through the present study, a uniform strategy of laboratory testing for CRS and population survey of pregnant women was established for the first time in India. The study reports the critical evaluation of the comparability between Euroimmun and Enzygnost ELISA kits for rubella specific IgM in suspected cases of congenital rubella syndrome and rubella specific IgG in serosurvey of pregnant women. Previous reports from India on CRS in different “at risk groups” (7), have used different testing strategies for laboratory confirmation of CRS.

Our results demonstrate satisfactory performance of both kits. Good individual reproducibility was observed, as evidenced by intra assay precision, with slightly higher CV for inter assay precision (Table 3&4). Mean absorbance and extinction ratio of positive control wells for both IgM and IgG were significantly higher in Euroimmun assays. Both the Euroimmun and Enzygnost kits are based on the principle of indirect ELISA, with purified antigen obtained from inactivated cell lysate of rubella virus, propagated in different cell lines. Contamination to some extent with cellular by-products, leading to non specific binding and reactions is therefore expected (8). By using a control antigen well, in addition to a viral antigen well for each sample (9) as well as a correction factor calculated from anti rubella reference (mean of PP1 and PP2), the Enzygnost assay attempts to minimize nonspecific results. This would have contributed to lower extinction ratios for the Enzygnost assays.

Our data shows a good qualitative agreement between the two ELISA kits for detection of both anti rubella IgM and IgG (Table1). Sensitivity was observed to be 100% for Euroimmun IgM testing as compared to Enzygnost results, with specificity of 93.1%.This indicates that while no false negative results were identified by Euroimmun, there is a risk of false positivity, which was resolved by adopting a third testing strategy (CLIA). Considering that inter rater agreement, in this case inter kit agreement, may be due to chance, we have applied Cohen’s kappa statistics to understand the true agreement between the two kits. Our data (κ=0.86) shows “almost perfect agreement” as per Jacob Cohen’s original interpretation (10). Recent interpretation (11) identifies the κ to indicate strong agreement. In case of anti rubella IgG testing by Euroimmun, while specificity was 100%, indicating that no false positive results were obtained, risk of detection of false negative results remained with a sensitivity of 95.9%.Cohen’s kappa was 0.84, indicating almost perfect (10) or strong agreement (11) between the two tests.

Discordance was observed in 15 (5.3%) of IgM and 16 (3.7%) of IgG results. While it was not possible to test all these samples by an alternative method (CLIA), those tested could be classified as true positive or negative based on best of three test results. Some of the discordant results observed in our study might be the result of variations in the types and source of antigen used in the different assays

Quantitative comparison of IgG results by Bland Altman analysis was also satisfactory (Fig 1). Some variation in rubella specific IgG results using different ELISA kits is expected and widely recognized (12). Since the 1980s, rubella virus IgG assays have been calibrated against the same WHO international standard rubella virus serum (RUBI-1-94) (12). Much of the variation, stems from the difference between the nature of the analyte and tests originally applied for determining the standard and the currently used test systems (13).

Data from our study (Table1), shows that a slight overestimation of IgM positivity(~5%) is possible with Euroimmun ELISA. The issue could be resolved by correlation of reports of individual suspected CRS cases with clinical presentation and other investigational findings.

Resolution of IgG testing and interpretation is usually required for serosurvey in an adult population, where high population prevalence is expected. There is a possibility of reporting false negative IgG results using the Euroimmun kit, as noted in our study (~3%). An earlier evaluation of eight commercial rubella IgG assays, also had a similar observation(14). Reporting a false negative rubella specific IgG result is considered more acceptable than a false-positive result. Such reports can possibly lead to vaccination or anxiety for a pregnant woman (14). However, a false positive report could result in missing a candidate for vaccination, who could get infected if exposed to the virus. The outcome of testing in CRS cases where sustained positivity of two sequential IgG tests are required for confirming the diagnosis, is beyond the scope of this study.

In conclusion our results show good agreement of Euroimmun and Enzygnost kits for rubella specific IgM (qualitative) and IgG (qualitative and quantitative). Euroimmun ELISA may be considered for detection of rubella specific IgM in suspected CRS cases and rubella specific IgG in surveillance studies.

## Acknowledgements

Dr JP Muliyil for critical review of manuscript.

## Funding Agency

UNDP through GAVI HSS and ICMR

## Author Contribution

Planned the study: *GNS,MVM,NG,RV,AMA;* Centre wise testing and interpretation: *AMA, MPS, VN, CR, AM, SN and MG;* CLIA analysis: *AMA,MPS;* Data compilation and analysis: *RV, SG, VJ;* QC: *VJ,SG and OK;* Statistical support: *SMJ;* First draft: *RV;* Critical review and final inputs: *GNS*. All authors read and approved the final draft of the manuscript

